# Context-explorer: Analysis of spatially organized protein expression in high-throughput screens

**DOI:** 10.1101/374751

**Authors:** Joel Ostblom, Emanuel J.P. Nazareth, Mukul Tewary, Peter W. Zandstra

**Author notes:** Corresponding Author: Peter W. Zandstra, Professor, Canada Research Chair in Stem Cell Bioengineering, Institute of Biomaterials and Biomedical Engineering, Vancouver Campus, 2185 East Mall, Vancouver, BC Canada V6T 1Z4.

## Abstract

A growing body of evidence highlights the importance of the cellular microenvironment as a regulator of phenotypic and functional cellular responses to perturbations. We have previously developed cell patterning techniques to control population context parameters, and here we demonstrate Context-explorer (CE), a software tool to improve investigation of microenvironmental variables through colony level analyses. We demonstrate the capabilities of CE in the analysis of human and mouse pluripotent stem cells (hPSCs, mPSCs) patterned in colonies of defined size and shape in multi-well plates.

CE employs a density-based clustering algorithm to identify cell colonies within micropatterned wells. Using this automatic colony classification methodology, we obtain accuracies comparable to manual colony counts in a fraction of the time. Classifying cells according to their relative position within a colony enables statistical analysis of radial spatial trends in protein expression within multiple colonies in the same treatment group. When applied to colonies of hPSCs, our analysis reveals a radial gradient in the expression of the pluripotency inducing transcription factors SOX2 and OCT4, and a similar trend in the intra-colony location of different cellular phenotypes. We extend these analyses to colonies of different sizes and shapes and demonstrate how the metrics derived by CE can be used to asses the patterning fidelity of micropatterned plates.

We have incorporated a number of features to enhance the usability and utility of CE. To appeal to a broad scientific community, all of the software’s functionality is accessible from a graphical user interface, and convenience functions for several common data operations are included. CE is compatible with existing image analysis programs such as CellProfiler and extends the analytical capabilities already provided by these tools. Taken together, CE facilitates investigation of spatially heterogeneous cell populations in fundamental research and drug development validation programs.

## Introduction

Emerging pieces of evidence stress the importance of a cell’s local microenvironment as a regulator of cellular phenotype and gene expression heterogeneity within cell populations. Microenvironmental parameters such as mechanical forces, cell to cell contact and endogenous signaling, all vary between cells at different positions in a well (McBeath et al., 2004, Peerani et al., 2007). The spatial heterogeneity of these factors leads to variability in efficiency of endocytosis and the vulnerability to viral infection (Snijder et al., 2009), influences epithelial tissue growth (Kim et al., 2009), impacts the expression of angiogenic factors in tumor cells (Kumar et al., 1998) and influences the differentiation potential of mouse and human pluripotent stem cells (mPSCs, hPSCs) (Davey and Zandstra, 2006, Peerani et al., 2007). Microenvironmental heterogeneity is also a potential confounding factor behind contradictory findings in the response of different cell types to key signaling pathway activity (Akopian et al., 2010, Jong et al., 2009, Snijder et al., 2012) and could limit the interpretation and reproducibility of experiments.

A comparative analysis (Haibe-Kains et al., 2013) of two large scale pharmacogenomic studies, the Cancer Genome Project (Garnett et al., 2012) and Cancer Cell line Encyclopedia (Barretina et al., 2012), revealed a surprisingly poor correlation between cell line drug response phenotypes between laboratories, which prevented meaningful extraction of drug-gene relationships. Correlation remained low even when using matched protocols and cell lines with highly correlated gene expression profiles. Although the exact source of variation in this study is unknown, a separate analysis of single cell data from 45 high-throughput (HTP) screens revealed that population context is indeed a ubiquitous source of variation between screens, and accounting for population context can improve experimental reproducibility between cell lines and laboratories (Snijder et al., 2012). Although it is acknowledged that understanding population heterogeneity is critical in biomedical research (Altschuler and Wu, 2010, Pelkmans, 2012), the scientific community has been slow to adopt approaches to reduce heterogeneity, such as controlling microenvironmental variables.

The increasing affordability of high content screening instruments, emergence of core screening facilities and technological advancements such as micropatterning in multi-well plates (Azioune et al., 2009), enable investigation of population context dependent variables with unprecedented throughput and veracity (Xia and Wong, 2012). By patterning extracellular matrix (ECM) proteins on a tissue culture surface, cells can be restricted to adhere to an array of spots of predefined shapes and sizes (Fink et al., 2007, Folch et al., 2000). An advantage of such patterning is enhanced control over microenvironmental variation within each well and improved assay robustness (Nazareth et al., 2013, @nazareth_multi-lineage_2016). Growing cells in colonies of defined size and shape facilitates analyses of inter- and intra-colony variation in protein expression. For example, we observe that hPSCs growing in such patterned colonies express varying levels of pluripotency markers OCT4 and SOX2 depending on colony size (Nazareth et al., 2013, Peerani et al., 2007). Elucidating the impact of the population context dependent variables on cellular phenotype will not only add to our understanding of fundamental cell biology, but will also allow us to optimize culture conditions and cell assays, provide possible explanations for current seemingly conflicting research and inform in silico models. These aspects are critical to next-generation drug development strategies and systems biology approaches.

We have previously developed an HTP platform for micropatterning of cells on ECM spots of defined shapes in multi-well plates (Nazareth et al., 2013, Tewary et al., 2017). To augment this platform, we here present a computational tool, Context-explorer (CE), which facilitates colony level analyses and cell patterning quality control. The CE software is meant to extend the functionality of currently available software solutions, both open source and commercial, for analyzing features of imaged cells (Carpenter et al., 2006, Jones et al., 2008, Misselwitz et al., 2010). While some implementations already exist that can be used to identify arrays of cells on glass slides (Bauer et al., 2012) or to study differential gene expression of cells in different spatial locations within the same colony (Gorman et al., 2014), our software aims to improve the HTP workflow for analysing cells in micropatterned multi-well plates by facilitating evaluation of patterning fidelity, enabling identification of colonies within a well, and improving spatial analyses of heterogeneous protein expression within colonies. As HTP technologies become more widespread, it is increasingly important to provide user friendly data analysis software targeted towards these platforms. Here, we demonstrate the utility of CE by investigating the impact of intra-colony location on hPSC pluripotency marker expression.

### Design and Implementation

Designed to complement existing imaging software, CE fits into the analysis pipeline following the extraction of cellular features from microscope images (**Fig 1A**). The input to CE is a CSV-file, which contains single cell xy-coordinates and at least one other measurement of interest, such as protein fluorescent intensity values. These single cell coordinates can be clustered into colonies within which spatial trends for the measurements of interest can be visualized (**Fig 1B**). By leveraging existing image extraction software and processing the resulting text files, CE has low system requirements and runs smoothly on modern laptop computers. CE is implemented in Python, and utilizes the scientific open source ecosystem SciPy (Jones et al., 2001). Specifically, NumPy (Walt et al., 2011) and Pandas (McKinney, 2010) are used for array manipulations, while Matplotlib (Hunter, 2007) and Seaborn (Waskom et al., 2016) generate the graphical visualizations. To make CE easily accessible to a broad scientific community of various technical backgrounds, all functionality is available via a graphical user interface designed in Qt.

**Figure 1:**
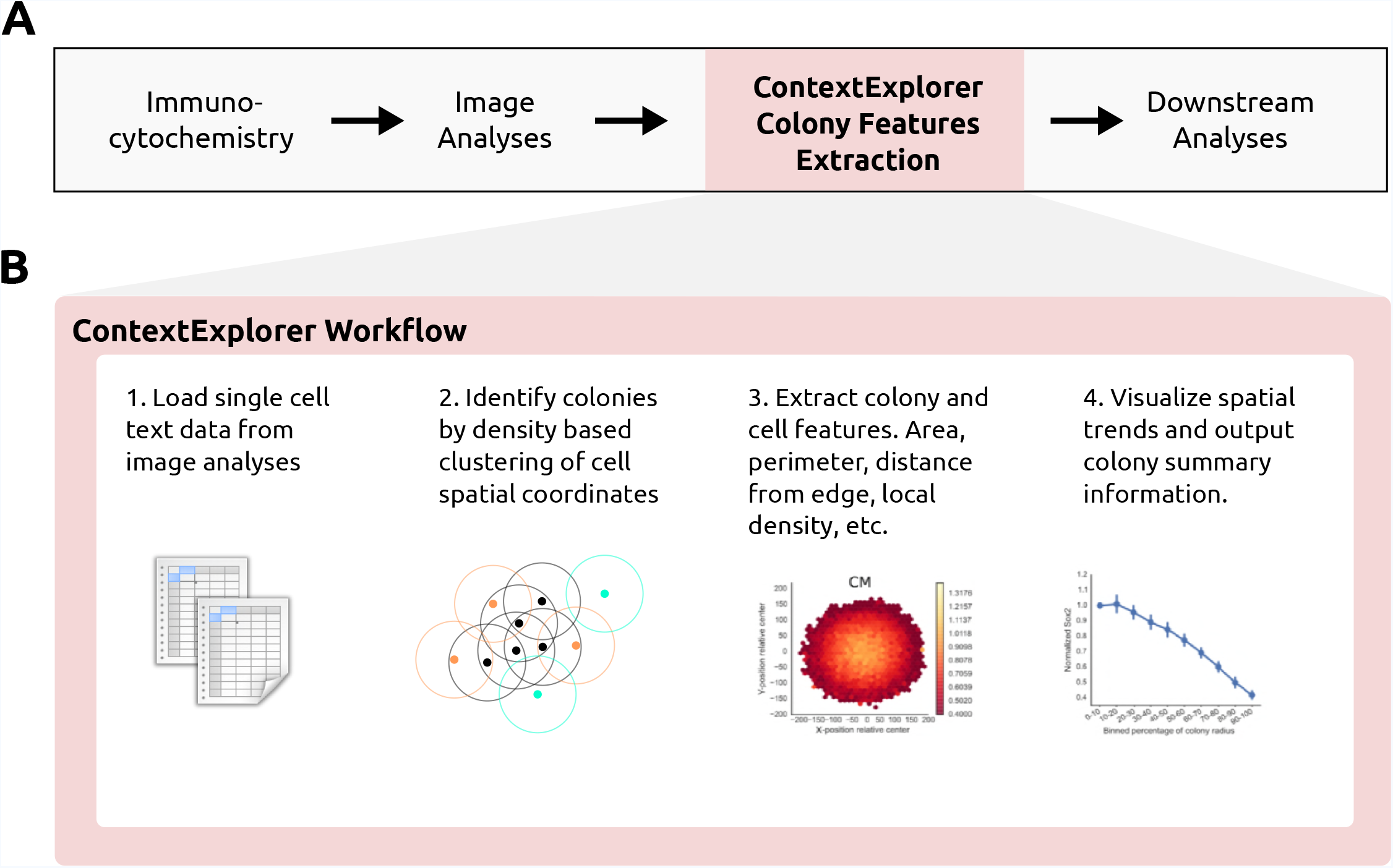
CE context and workflow in the image analysis pipeline. **A)** The input to CE is a CSV-file with single cell measurements of interest. CE thus fits into existing image analysis pipelines after these measurements have already been extracted from the images. **B)** The workflow of using CE, each step is described in detail in the methods section.

To interrogate organized cell behavior within colonies, the concept of cellular colonies must first be introduced by classifying closely positioned cells as belonging to the same colony. Manually labeling individual cells is infeasible in HTP assays that often include millions of cells. There are many existing algorithms for automatically identifying dense clusters of data points (Rui Xu and Wunsch, 2010) and CE employs the Density-Based Spatial Clustering of Applications with Noise (DBSCAN) algorithm (Ester et al., 1996), as implemented in the scikit-learn Python package (Pedregosa et al., 2011), to identify sets of points at high two dimensional density. Clustering cells into colonies based on local cell density is similar to how these groups are defined biologically since cellular communication is restricted by the distance between cells. DBSCAN scan also has the advantage that it has a notion of outliers, cells far away from any colony, and can classify such cells as noise rather than trying to force all cells to belong to a colony as other cluster algorithms. DBSCAN performs unsupervised clustering and does not require prior knowledge of the number of colonies within each well, only specification of the neighborhood search radius (Eps) and the minimum number of points (MinPts) within the neighborhood to start propagating a cluster. For each point found within the Eps neighbourhood of the starting point, a search for additional points will be performed. If the number of points found within a point’s Eps neighborhood is greater or equal to MinPts, that point is considered a core point of the cluster. Points that fail to meet this criteria, but that are density reachable from a core point, are considered border points and are classified as part of the colony. Points that fail to meet either of these two criteria are labelled as noise and not part of any colony.

While there are implementations of DBSCAN that automate parameter optimization, these increase time complexity (Ankerst et al., 1999, Karami and Johansson, 2014). As an alternative to automatic parameter estimation, CE allows for the Eps and MinPts parameters to be adjusted via the graphical user interface while viewing the resulting colony identification accuracy. The immediate visual feedback enables intuitive and accurate colony classification and decreases the time it takes to optimize Eps and MinPts. DBSCAN clustering is deterministic for the core cells of each cluster and only border points which are density reachable from more than one cluster core can be assigned to different clusters between runs. Colonies in ECM patterned wells rarely grow close enough for border cells to be density reachable from more than one colony, so for this application cells are routinely clustered deterministically. To further increase colony identification accuracy, CE includes filters for colony size, density and circularity, which refine the colony identification procedure and are particularly useful to deal with imaging artefacts and overgrown colonies.

Each cluster of points returned by DBSCAN corresponds to cells growing together in a colony. To define colony attributes, CE utilizes the geometric analyses package Shapely (Gillies and others, 2007). Generally, the polygonal boundary area of a colony can be defined as either the convex or concave hull of its cells. CE uses the convex hull algorithm as it is less expensive to compute than the concave hull and performs well with commonly used micropatterned spot shapes. By finding the colony bounding area, additional geometric attributes such as colony diameter, circumference, area and cell density can be calculated for each colony. Additionally, each cell can be assigned Cartesian relative the colony’s centroid or the closest edge of the colony’s boundary area. These relative cell coordinates can be used to group cells at similar positions from multiple colonies. The process for grouping cells is initiated by deriving an aggregated value of all the cells in a colony that are within the same location bin. These colony values are then aggregated for multiple colonies and the error estimation reported describes the variation between colonies rather than between cells within one colony. The visualizations built into CE further facilitates the analyses of spatial trends within colonies. Cells can also be grouped according to a hexagonal grid, which aggregates cells from all colonies in the same bin within the grid.

## Results and Discussion

### Colony classification

To demonstrate the colony classification process, we patterned mouse PSCs on circular ECM spots 500 um in diameter in a 96-well plate using an in-house HTP UV-lithography method (Tewary et al., 2017). To enable extraction of cellular coordinates within the well, cell nuclei were labelled with DAPI and analysed by a primary image analysis program, such as CellProfiler. After imaging and extraction of single cell features, the resulting CSV-file was processed by CE to classify cells into colonies. Cells robustly adhered to the patterned regions, and were grown for 48 h in pluripotent conditions (LIF and Serum). These micropatterned, well-separated clusters of cells were easily identified after adjusting the DBSCAN Eps parameter so that the distance between colonies in a micropatterned well is longer than the threshold for the intercellular distance within the same colony. However, cells occasionally bridge adjacent confluent colonies, effectively merging them together. Such colonies would still be classified as valid clusters by DBSCAN, since all the cells are density reachable from each other. Another important caveat is that the imaging hardware may not allow the entire well to be captured, resulting in partial images of many of the colonies. Including either of these merged or partial colonies in the downstream analyses, could confound the interpretation of the underlying biology.

When limited to only the default DBSCAN parameters MinPts and Eps, partial and merged colonies are difficult to discriminate from colonies of desired shape and size. We found that an efficient way to eliminate these undesired colonies, was to apply a set of filtering criteria to the colonies detected by DBSCAN. Filtering on colony roundness and size were the most effective criteria to exclude merged and partial colonies from the DBSCAN output (**Fig 2A**). The effect of applying size and roundness filters was striking when comparing the number of cells per colony before and after filtering. Prior to filtering, several clusters of colony sizes were detected, including bigger merged colonies and smaller partial colonies cut off by the imaging limitations (**Fig 2B**). These colonies were omitted from the final analyses as they would skew the calculations of both the mean number of cells per colony and spatial trends within the colonies. After excluding colonies of undesirable size and shape, we observed only one cluster of colony sizes and the mean number of cells per colonies was notably consistent between wells, indicating reproducible patterning of cells (**Fig 2C**).

**Figure 2:**
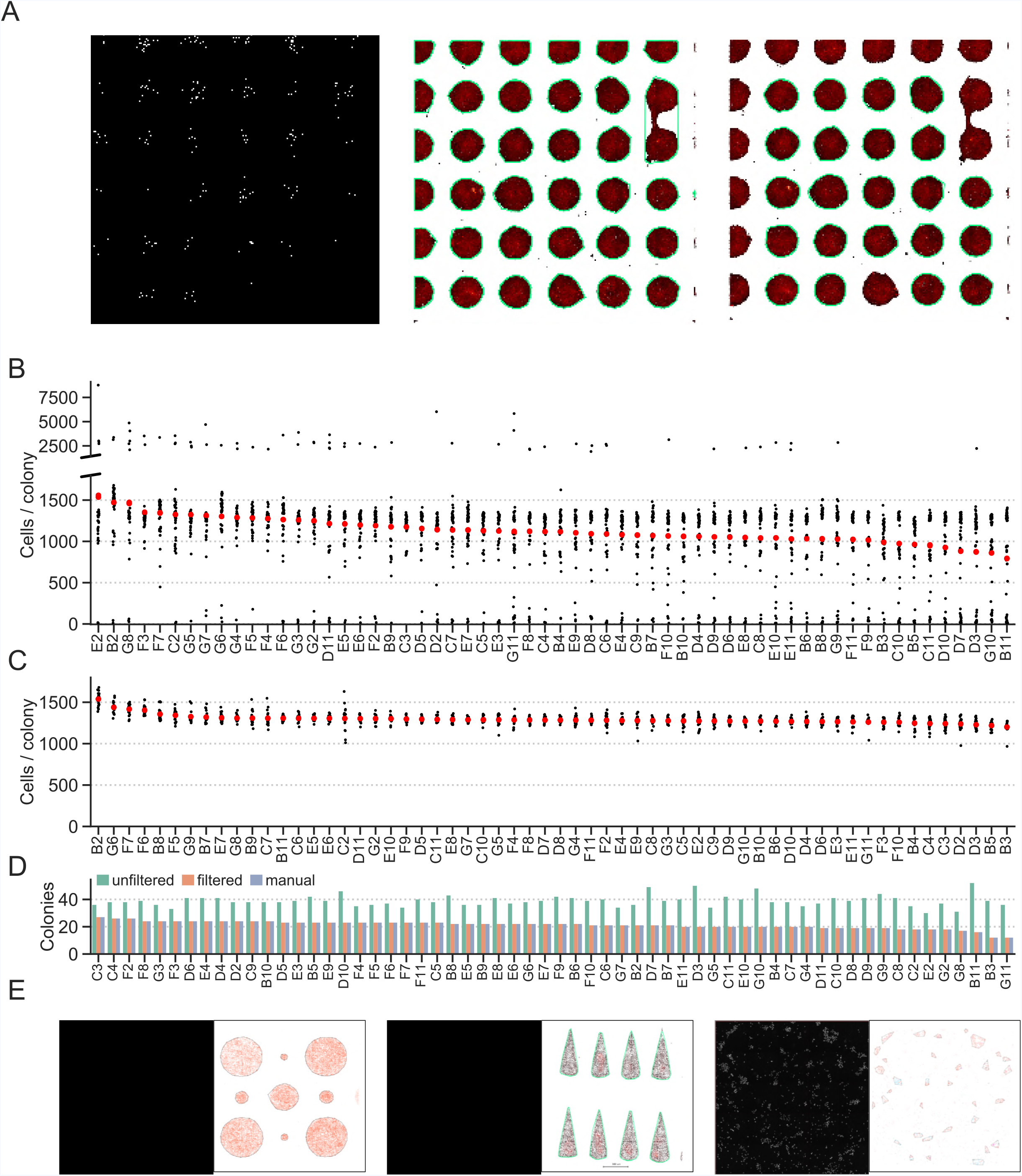
Classification of cells into colonies. **A)** mPSCs restricted to grow on micropatterned ECM spots (*left*) can be classified into colonies via the DBSCAN algorithm (*middle*). Merged and partial colonies can be excluded by applying filters to the DBSCAN clustering results (*right*). Each data point represents a cell and colonies are encircled with green lines. **B)** Number of cells per colony in wells from a 96-well plate after running the DBSCAN. **C)** Number of cells per colony in wells after excluding merged and partial colonies from the DBSCAN results. **D**) Number of colonies per well after running DB-SCAN with and without filters, and from a visual inspection of the images. **E**) Colony identification for colonies of different sizes in the same well (*left*), colonies of non-circular geometries (*middle*), and colonies in wells without pre-patterned ECM spots (*right*).

Another useful metric for assessing patterning fidelity is the number of colonies per well, which is also computed by CE. Comparing the number of filtered and unfiltered colonies identifies wells containing many fused and partial colonies that were detected by DBSCAN, but then excluded by the filters. The filtered count was nearly identical to the counts obtained from visually inspecting the images from each well (**Fig 2D**), and were computed in a fraction of the time of manual counting. CE can also accurately identify clusters of cells growing in micro-patterned ECM spots of different sizes within the same well, cells patterned with UV-lithography in non-circular shapes, and in unpatterned wells (**Fig 2E**). These results demonstrate the capacity for CE to identify colonies of cells in ECM patterned wells with an accuracy similar to that of visual image inspection.

### Investigating the behavior of hPSCs in micropatterned colonies

To apply CE to hPSCs analysis, we first patterned hPSCs in 200 μm diameter colonies in 96-well plates using microcontact-printing. While most cells adhere to the ECM spots in the patterned plates, there is also limited non-specific cell adhesion in-between patterned ECM spots. Compared to UV-lithography, microcontact printing has a higher proportion of cells growing in tiny colonies and as single cells outside patterned ECM spots, which makes this technology suitable for comparing the behavior of cells outside and inside micropatterned colonies. To test whether cells that adhere non-specifically display differences in protein expression compared to cells within colonies, we assessed cellular response 42 h after treatment with serum free medium containing BMP4 (SF+B, induces trophectoderm and primitive endoderm (Vallier et al., 2009, Xu et al., 2002)), or MEF conditioned medium (CM, maintains pluripotency (Xu et al., 2001)). Expression of the pluripotency-associated transcription factor SOX2 was analyzed to quantify cellular differences. As expected, in SF+B medium, pluripotency signals were repressed and no difference was observed in SOX2 expression between cells inside and outside colonies. In contrast, CM induces the expression of SOX2 in cells within colonies, while cells outside colonies express the marker to a lesser extent (**Fig 3A**). This is visible both in a change of the distribution shape and a shift in means of the SOX2 intensities. These differences in absolute fluorescence intensities suggest that cells inside and outside colonies do not respond similarly to added factors in the medium, further highlighting the importance of controlling for microenvironmental parameters such as population context when assessing cellular responses to experimental conditions.

**Figure 3:**
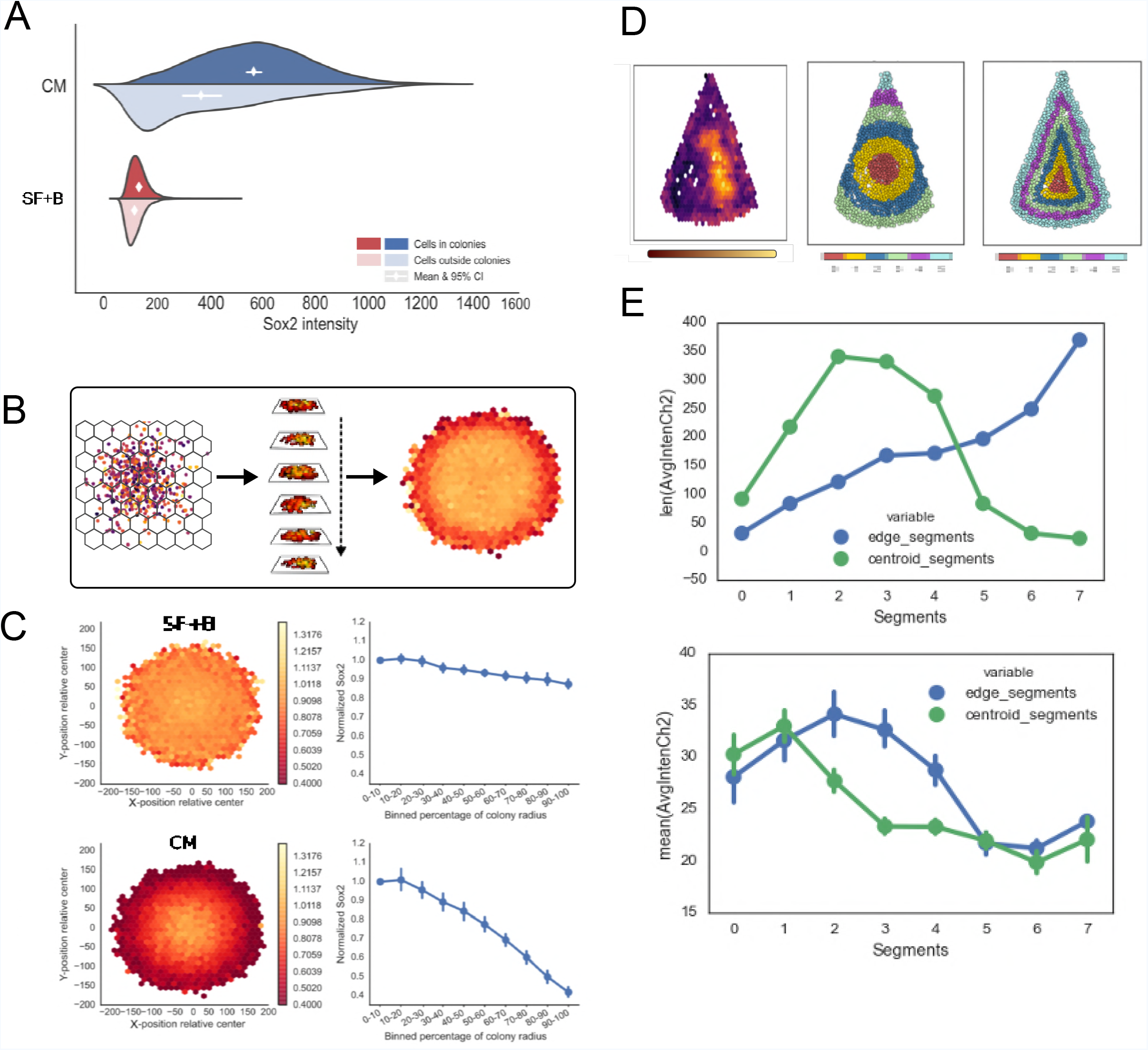
Quantification of radial expression trends. **A)** Comparison of SOX2 expression between cells growing inside and outside of colonies in microcontact-printed wells. **B)** Cells from multiple colonies at similar locations within their respective colony are aggregated together in bins according to a hexagonal grid system. The heatmap is colored by the desired measure of variation or central tendency. **C)** Multiple colonies of hPSCs are aggregated to reveal general tendencies in the spatial protein expression pattern of SOX2 (*left*). Each bin shows the mean expression of cells from multiple colonies. Trends are visualized as line plots, where joint data points represent the mean intensity of cells at each distance bin throughout the colony (*right*). Error bars represent 95% confidence intervals between cells from the same location in different colonies. **D)** SOX2 expression of triangular hPSC colonies (*left*). Cells grouped in radial segments according to their distance from the colony centroid (*middle*) or to the closest colony edge (*right*). **E)** The number of cells assigned to each bin with the binning techniques (distance from edge or centroid) in panel D (*top*). The average intensity values for the different bin segmentations (*bottom*).

### Analysis of spatial trends in protein expression within hPSC colonies

In addition to facilitating inter-colony analyses, CE allows for investigation of intra-colony variation in protein expression. Radial gradients of protein expression have previously been reported in non-patterned and patterned colonies of hPSCs (Davey and Zandstra, 2006, Peerani et al., 2007, Warmflash et al., 2014). Analysing these trends through visual inspection and manual data analyses is feasible in a low throughput platform, but becomes error-prone and time-consuming in HTP systems with hundreds or thousands of colonies. To visualize spatially biased protein expression, CE can automatically aggregate colonies within replicate wells and display a heatmap of spatial protein expression variation within these colonies (**Fig 3B**). This analysis can be applied to colonies of various sizes and shapes.

When investigating colonies of hPSCs grown in either CM or SF+B, we observed distinct radial gradients of SOX2 expression between the two conditions (**Fig 3C**). To distinguish differences attributed to spatial factors from those attributed to exogenous factors, all intensities were normalized relative to the expression at the centroid of the colony. In hPSCs grown in CM, SOX2-expression decreased in a linear fashion towards the edge of the colony, with cells at the colony border only displaying half the fluorescence intensity of cells near the colony centroid. Meanwhile, cells grown in SF+B, exhibited low SOX2 expression throughout the entire colony, which can be attributed to BMP4 inducing differentiation and overriding any local pluripotency supporting signals.

Quantitative evaluation of protein expression levels relative to the location of a cell within a colony was performed by aggregating cells in radial bins according to their distance from the colony centroid rather than their relative xy-coordinates. The mean or median expression values could then be compared between ring-shaped bins across multiple colonies. This analysis technique highlights the spatial trends of SOX2-expression for hPSCs grown in CM (**Fig 3C**), where expression of SOX2 was highest in cells at the center of the colony and linearly decreased toward the colony edge. Statistical significance at p=0.01 can be roughly inferred from non-overlapping pairs of 95% confidence intervals (Cumming and Finch, 2005). However, it should be noted that statistical significance is easily achieved with sample sizes this large, even at small effect sizes (Haney et al., 2014), so it is important to assess the magnitude of the differences. In hPSCs cultured in SF+B, a weak radial gradient of SOX2 expression emerged exhibiting no more than a 10% difference in expression level between cells at the centroid and the edge of the colony. In contrast, hPSCs cultured in CM exhibit more than double the level of SOX2 expression at the center of the colony compared to the edge.

For cells grown on ECM patterns of non-circular geometries, we evaluated how grouping cells into bins based either on the distance from the colony border or the colony centroid affected our interpretation of spatial trends in protein expression. To illustrate the different biological interpretations that could arise from these two metrics, we investigated the SOX2 expression of hPSC colonies grown on triangular ECM patterns in SF+B. Colonies were aggregated as described previously, and spatial bias in SOX2 expression was visualized (**Fig 3D**). To quantify these trends, colonies were grouped into concentric annular or triangular segments, based on their distance from the colony centroid or the colony border, respectively. Depending on the binning strategy, cells were grouped to different segments and the number of cells in each segment also differed greatly. Importantly, the choice of binning metric influences the interpretation of the resulting protein expression trends. In this example, SOX2 expression decreased more rapidly as a function of the distance from the colony centroid compared to from the colony edge (**Fig 3E**).

### Conclusions

There is overwhelming evidence that increased control and monitoring of population context parameters is needed to improve assay reproducibility and to understand heterogeneous responses between cells in the same population. However, addressing this challenge has proven difficult in the broader biomedical community. To augment the power of HTP analysis of population context parameters in the cellular microenvironment, we previously developed cell patterning techniques to control population context parameters, and here we demonstrate a software tool for improved monitoring of microenvironment variables in HTP assays. In this study, CE was utilized to explore and quantify radial spatial trends in SOX2 expression within micropatterned hPSC colonies of various shapes and sizes. We observed that the cellular phenotype varies as a function of location within a colony, further highlighting the importance of understanding variation in population context dependent factors.

### Availability and Future Directions

CE is compatible with existing HTP imaging software and standard fluorescent microscopy based assays. By developing a GUI-driven workflow and releasing it under an open source license, we provide a solution to facilitate colony-level analysis for a wide scientific community. Members of our group regularly use CE for colony level analyses as evidenced in previously published and ongoing studies. (Nazareth et al., 2013, 2016, Rahman et al., 2017, Tewary et al., 2017). To further broaden the utility and applications of the software, there are built-in visualizations to assist with fluorescent intensity thresholding and pattern fidelity assessment, and interface components to assign wells to treatment groups. To lower the threshold for wide adoption, CE is distributed as a binary package through the conda package manager, which runs under Linux, OS X and Windows. The source code is distributed under the 3-clause BSD open source license, which enables incorporation of its features into existing image analysis pipelines. Source code and installation instructions are available online at *https://gitlab.com/stemcellbioengineering/context-explorer*, and documentation is maintained at *https://contextexplorer.readthedocs.io*.

## Author Contributions

J.O. developed the software, analyzed the data, and performed the UV-lithography experiments for mPSCs. E.J.P.N. designed and performed the microcontact printing experiments. M.T. designed and performed the UV-lithography experiments for hPSCs. J.O., E.J.P.N. and P.W.Z. designed the project and wrote the manuscript.

## Acknowledgements

J.O is grateful to Gålöstiftelsen and NSERC M3 for supporting his work and we thank Céline Bauwens for critical reading and editing of the manuscript.

## Supplementary Methods

### Cell culture

We obtained the H9 hESC line (WA09) from the WiCell Research Institute. H9 cells were routinely cultured on feeder layers of irradiated murine embryonic fibroblast (MEF) feeders in knockout (KO)-Dulbecco’s modified Eagle’s medium (DMEM) (Invitrogen) with 20% KO-serum replacement (Invitrogen) (KO-DMEM) supplemented with 4 ng mL–1 FGF-2 (PeproTech). Cells were passaged at 1:4 to 1:6 split ratios every 4-5 days by dissociating colonies with 0.1% collagenase IV (Invitrogen) into small clumps. All cell line stocks were confirmed negative for mycoplasma contamination.

### Micropatterning of cells into 96-well plates

#### Microcontact printing

We previously developed a method for patterning proteins in standard 96-well plates using microcontact printing (Nazareth et al., 2013). The PDMS stamps were fabricated using standard soft lithography techniques, with the exception that liquid PDMS was cast into a Teflon mould before curing, allowing control of the shape of the PDMS stamp. Microcontact printing was carried out according to our previously published protocol (Peerani et al., 2009). Briefly, the ECM solution (Matrigel diluted 1:30 in phosphate buffered saline) was deposited onto the patterned surface of ethanol sterilized PDMS stamps for 4 h at room temperature. Stamps were rinsed with ddH2O, dried gently with N2 gas, placed into tissue-culture treated 96-well plates, and incubated in the 96-well plates for 7-10 min in a humidity chamber (Relative humidity 55-70%). The stamps were then removed and substrates were passivated with 5% weight Pluronic F-127 (Sigma-Aldrich) in ddH20 for 1 h.

#### UV lithography

We recently developed an alternative method to microcontact printing using UV-lithography to pattern proteins in standard 96-well (Tewary et al., 2017). Briefly, glass cover-slips were activated in a plasma cleaner and rinsed in ddH2O. Patterns of predefined size and shape were transferred to the cover-slip by photo-oxidizing select regions of the substrate using Deep UV exposure. The patterned slides were assembled to bottomless 96-well plates to produce plates with patterned cell culture surfaces. Prior to seeding cells onto the plates, the wells were activated with N-(3-Dimethylaminopropyl)-N-ethylcarbodiimide hydrochloride (Sigma cat# 03450), and N-Hydroxysuccinimide (Sigma, cat# 130672) for 20 minutes. Before seeding cells, the plates were incubated with Geltrex (for hPSC) or 12.5 ug / ml fibronectin in gelatin (for mPSCs) and washed with ddH2O.

### Seeding hPSCs onto patterned substrates

The hPSCs were dissociated using TrypLE™ for three min. TrypLE™ was inactivated by adding medium containing 20% KO-SR. Cells were centrifuged and resuspended in Nutristem^®^ hESC XF (Biological Industries, cat. no. 05-100-1A) and 10 μM ROCK inhibitor Y-27632 (Tocris). The SF medium contains DMEM/F12, 1x Nonessential amino acids, 50 U/mL Penicillin, 50 μg/mL Streptomycin, 10 μg/mL bovine Transferrin, 0.1 mM ß-Mercaptoethanol (all Invitrogen), 2% fatty acid-free Cohn’s fraction V BSA (Sero-logicals), 1x Trace Elements A, B & C (Mediatech), 50 μg/mL Ascorbic Acid (Sigma) and 7 μg/mL recombinant human insulin. Cells were seeded at 105 cells per well (or as described in text) and incubated. After 6 h, cells were washed twice with PBS and incubated for another 42 h in fresh medium (either SF supplemented with factors or CM).

### Immunocytochemistry

Plates were fixed for 30 min in 3.7% formaldehyde and permeabilized for 3 min in 100% methanol. Next the plates were incubated overnight at 4°C with primary antibodies in 10% FBS in PBS. Finally, the plates were incubated with AlexaFluor secondary antibodies (1:200; Molecular Probes) for 1 h at room temperature in 10% FBS in PBS. Fluorescent images were obtained for expression of OCT4 (1:200 BD), SOX2 (1:200 R&D Systems), and Hoechst 33342 (Sigma-Aldrich).

### High-content image analysis

Plates were imaged and analyzed using the Cellomics Arrayscan VTI platform and Target Activation protocol (Thermo Scientific). This protocol generates nuclear masks, provides single cell average nuclear intensity values for protein expression and DNA content, as well as spatial x- and y-coordinates of the nuclei centroids. Single cell x-y- coordinates and fluorescent intensity data were exported as CSV-files and imported into CE for exploration of colony level details.

### Statistical analysis

Error bars on plots represent 95% confidence intervals (CI) of replicates except where indicated differently. CIs are calculated by resampling the original distribution 1000 times as implemented in Seaborn.

## References

Akopian, V., Andrews, P.W., Beil, S., Benvenisty, N., Brehm, J., Christie, M., Ford, A., Fox, V., Gokhale, P.J., Healy, L., et al. (2010). Comparison of defined culture systems for feeder cell free propagation of human embryonic stem cells. In Vitro Cellular & Developmental Biology - Animal 46, 247–258.

Altschuler, S.J., and Wu, L.F. (2010). Cellular Heterogeneity: Do Differences Make a Difference? Cell 141, 559–563.

Ankerst, M., Breunig, M.M., Kriegel, H.-p., and Sander, J. (1999). OPTICS: Ordering Points To Identify the Clustering Structure. (ACM Press), pp. 49–60.

Azioune, A., Storch, M., Bornens, M., Théry, M., and Piel, M. (2009). Simple and rapid process for single cell micro-patterning. Lab on a Chip 9, 1640–1642.

Barretina, J., Caponigro, G., Stransky, N., Venkatesan, K., Margolin, A.A., Kim, S., Wilson, C.J., Lehár, J., Kryukov, G.V., Sonkin, D., et al. (2012). The Cancer Cell Line Encyclopedia enables predictive modelling of anticancer drug sensitivity. Nature 483, 603–607.

Bauer, M., Kim, K., Qiu, Y., Calpe, B., Khademhosseini, A., Liao, R., and Wheeldon, I. (2012). Spot Identification and Quality Control in Cell-Based Microarrays. ACS Combinatorial Science 14, 471–477.

Carpenter, A.E., Jones, T.R., Lamprecht, M.R., Clarke, C., Kang, I.H., Friman, O., Guertin, D.A., Chang, J.H., Lindquist, R.A., Moffat, J., et al. (2006). CellProfiler: Image analysis software for identifying and quantifying cell phenotypes. Genome Biology 7, R100.

Cumming, G., and Finch, S. (2005). Inference by Eye: Confidence Intervals and How to Read Pictures of Data. American Psychologist 60, 170–180.

Davey, R.E., and Zandstra, P.W. (2006). Spatial Organization of Embryonic Stem Cell Responsiveness to Autocrine Gp130 Ligands Reveals an Autoregulatory Stem Cell Niche. STEM CELLS 24, 2538–2548.

Ester, M., Kriegel, H.-p., S, J., and Xu, X. (1996). A density-based algorithm for discovering clusters in large spatial databases with noise. (AAAI Press), pp. 226–231.

Fink, J., Théry, M., Azioune, A., Dupont, R., Chatelain, F., Bornens, M., and Piel, M. (2007). Comparative study and improvement of current cell micro-patterning techniques. Lab on a Chip 7, 672–680.

Folch, A., Jo, B.-H., Hurtado, O., Beebe, D.J., and Toner, M. (2000). Microfabricated elastomeric stencils for micropatterning cell cultures. Journal of Biomedical Materials Research 52, 346–353.

Garnett, M.J., Edelman, E.J., Heidorn, S.J., Greenman, C.D., Dastur, A., Lau, K.W., Greninger, P., Thompson, I.R., Luo, X., Soares, J., et al. (2012). Systematic identification of genomic markers of drug sensitivity in cancer cells. Nature 483, 570–575.

Gillies, S., and others (2007). Shapely: Manipulation and analysis of geometric objects (toblerity.org).

Gorman, B.R., Lu, J., Baccei, A., Lowry, N.C., Purvis, J.E., Mangoubi, R.S., and Lerou, P.H. (2014). Multi-Scale Imaging and Informatics Pipeline for In Situ Pluripotent Stem Cell Analysis. PLoS ONE 9, e116037.

Haibe-Kains, B., El-Hachem, N., Birkbak, N.J., Jin, A.C., Beck, A.H., Aerts, H.J.W.L., and Quackenbush, J. (2013). Inconsistency in large pharmacogenomic studies. Nature 504, 389–393.

Haney, S.A., Guey, L., and Chakravarty, A. (2014). Analyzing Cell-Level Data - An Introduction to High Content Screening. An Introduction to High Content Screening: Imaging Technology, Assay Development, and Data Analysis in Biology and Drug Discovery 145.

Hunter, J.D. (2007). Matplotlib: A 2D Graphics Environment. Computing in Science & Engineering 9, 90–95.

Jones, E., Oliphant, T., Peterson, P., and others (2001). SciPy: Open source scientific tools for Python.

Jones, T.R., Kang, I.H., Wheeler, D.B., Lindquist, R.A., Papallo, A., Sabatini, D.M., Golland, P., and Carpenter, A.E. (2008). CellProfiler Analyst: Data exploration and analysis software for complex image-based screens. BMC Bioinformatics 9, 482.

Jong, D. de, Koster, A., Hagenbeek, A., Raemaekers, J., Veldhuizen, D., Heisterkamp, S., Boer, J.P. de, and Glabbeke, M. van (2009). Impact of the tumor microenvironment on prognosis in follicular lymphoma is dependent on specific treatment protocols. Haematologica 94, 70–77.

Karami, A., and Johansson, R. (2014). Choosing DBSCAN Parameters Automatically using Differential Evolution. International Journal of Computer Applications 91, 1–11.

Kim, J.-H., Kushiro, K., Graham, N.A., and Asthagiri, A.R. (2009). Tunable interplay between epidermal growth factor and cell–cell contact governs the spatial dynamics of epithelial growth. Proceedings of the National Academy of Sciences 106, 11149–11153.

Kumar, R., Kuniyasu, H., Bucana, C.D., Wilson, M.R., and Fidler, I.J. (1998). Spatial and temporal expression of angiogenic molecules during tumor growth and progression. Oncology Research 10, 301–311.

McBeath, R., Pirone, D.M., Nelson, C.M., Bhadriraju, K., and Chen, C.S. (2004). Cell Shape, Cytoskeletal Tension, and RhoA Regulate Stem Cell Lineage Commitment. Developmental Cell 6, 483–495.

McKinney, W. (2010). Data Structures for Statistical Computing in Python. In Proceedings of the 9th Python in Science Conference, S. van der Walt, and J. Millman, eds. pp. 51–56.

Misselwitz, B., Strittmatter, G., Periaswamy, B., Schlumberger, M.C., Rout, S., Horvath, P., Kozak, K., and Hardt, W.-D. (2010). Enhanced CellClassifier: A multi-class classification tool for microscopy images. BMC Bioinformatics 11, 30.

Nazareth, E.J.P., Ostblom, J.E.E., Lücker, P.B., Shukla, S., Alvarez, M.M., Oh, S.K.W., Yin, T., and Zandstra, P.W. (2013). High-throughput fingerprinting of human pluripotent stem cell fate responses and lineage bias. Nature Methods 10, 1225–1231.

Nazareth, E.J.P., Rahman, N., Yin, T., and Zandstra, P.W. (2016). A Multi-Lineage Screen Reveals mTORC1 Inhibition Enhances Human Pluripotent Stem Cell Mesendoderm and Blood Progenitor Production. Stem Cell Reports 6, 679–691.

Pedregosa, F., Varoquaux, G., Gramfort, A., Michel, V., Thirion, B., Grisel, O., Blondel, M., Prettenhofer, P., Weiss, R., Dubourg, V., et al. (2011). Scikit-learn: Machine Learning in Python. Journal of Machine Learning Research 12, 2825–2830.

Peerani, R., Rao, B.M., Bauwens, C., Yin, T., Wood, G.A., Nagy, A., Kumacheva, E., and Zandstra, P.W. (2007). Niche-mediated control of human embryonic stem cell self-renewal and differentiation. The EMBO Journal 26, 4744–4755.

Peerani, R., Bauwens, C., Kumacheva, E., and Zandstra, P.W. (2009). Patterning mouse and human embryonic stem cells using micro-contact printing. Methods in Molecular Biology (Clifton, N.J.) 482, 21–33.

Pelkmans, L. (2012). Using Cell-to-Cell Variability—A New Era in Molecular Biology. Science 336, 425–426.

Rahman, N., Brauer, P.M., Ho, L., Usenko, T., Tewary, M., Zúñiga-Pflücker, J.C., and Zandstra, P.W. (2017). Engineering the haemogenic niche mitigates endogenous inhibitory signals and controls pluripotent stem cell-derived blood emergence. Nature Communications 8.

Rui Xu, and Wunsch, D.C. (2010). Clustering Algorithms in Biomedical Research: A Review. IEEE Reviews in Biomedical Engineering 3, 120–154.

Snijder, B., Sacher, R., Rämö, P., Damm, E.-M., Liberali, P., and Pelkmans, L. (2009). Population context determines cell-to-cell variability in endocytosis and virus infection. Nature 461, 520–523.

Snijder, B., Sacher, R., Rämö, P., Liberali, P., Mench, K., Wolfrum, N., Burleigh, L., Scott, C.C., Verheije, M.H., Mercer, J., et al. (2012). Single-cell analysis of population context advances RNAi screening at multiple levels. Molecular Systems Biology 8, 579.

Tewary, M., Ostblom, J., Prochazka, L., Zulueta-Coarasa, T., Shakiba, N., Fernandez-Gonzalez, R., and Zandstra, P.W. (2017). A stepwise model of Reaction-Diffusion and Positional-Information governs self-organized human peri-gastrulation-like patterning. Development dev.149658.

Vallier, L., Touboul, T., Chng, Z., Brimpari, M., Hannan, N., Millan, E., Smithers, L.E., Trotter, M., Rugg-Gunn, P., Weber, A., et al. (2009). Early cell fate decisions of human embryonic stem cells and mouse epiblast stem cells are controlled by the same signalling pathways. PloS One 4, e6082.

Walt, S. van der, Colbert, S.C., and Varoquaux, G. (2011). The NumPy Array: A Structure for Efficient Numerical Computation. Computing in Science & Engineering 13, 22–30.

Warmflash, A., Sorre, B., Etoc, F., Siggia, E.D., and Brivanlou, A.H. (2014). A method to recapitulate early embryonic spatial patterning in human embryonic stem cells. Nature Methods 11, 847–854.

Waskom, M., Botvinnik, O., drewokane, Hobson, P., Halchenko, Y., Lukauskas, S., War-menhoven, J., Cole, J.B., Hoyer, S., Vanderplas, J., et al. (2016). Seaborn: V0.7.0 (January 2016).

Xia, X., and Wong, S.T. (2012). Concise Review: A High-Content Screening Approach to Stem Cell Research and Drug Discovery. STEM CELLS 30, 1800–1807.

Xu, C., Inokuma, M.S., Denham, J., Golds, K., Kundu, P., Gold, J.D., and Carpenter, M.K. (2001). Feeder-free growth of undifferentiated human embryonic stem cells. Nature Biotechnology 19, 971–974.

Xu, R.-H., Chen, X., Li, D.S., Li, R., Addicks, G.C., Glennon, C., Zwaka, T.P., and Thomson, J.A. (2002). BMP4 initiates human embryonic stem cell differentiation to trophoblast. Nature Biotechnology 20, 1261–1264.

